# Substrate accessibility regulation of human TopIIa decatenation by cohesin

**DOI:** 10.1101/2023.11.20.567865

**Authors:** Erin E. Cutts, Sanjana Saravanan, Gemma L. M. Fisher, David S. Rueda, Luis Aragon

## Abstract

Human topoisomerase II alpha (TOP2α) is the main mitotic decatenase ^1–3^ resolving intertwines between sister chromatids that form during DNA replication ^4^. Here, we employ quadruple-trap optical tweezers to generate braids between a pair of λ-DNA molecules to study, at the single-molecule level and in real-time, how TOP2α untangles these DNAs. We find that TOP2α rapidly resolves single and multiple braids and is inhibited by chemotherapy agent, etoposide. TOP2α is sensitive to DNA conformation, exhibiting a chiral preference for the removal of braids with right-handed crossings and is inhibited when DNAs are held at forces over 20pN. We show that TOP2α must load at the cross between the two DNAs to resolve braids efficiently. TOP2α pre-loaded onto individual DNAs is unable to resolve newly formed braids in the presence of ATP, suggesting that the geometry of pre-bound enzyme is incompatible with the capture of the second DNA. Finally, we show that cohesin binds to braids, preventing TOP2α from resolving them. Our study unveils novel insights into the regulation of TOP2α’s decatenation, underscoring the importance of substrate accessibility and the role of cohesin in modulating TOP2α activity.

## Introduction

The double helical structure of DNA, consisting of two strands of nucleotides wound around each other in a spiral staircase-like manner ^5^, provides stability to the molecule, protecting the genetic information it carries. However, this structure also represents significant challenges during various cellular processes. One such obstacle is the topological constrains arising from the twisting and winding of the DNA molecule. For example, as DNA is unwound during transcription, torsional strain is introduced on the molecule generating supercoiling ^6^. Similarly, as replication forks progress and the parental DNA strands separate, intertwines or catenanes are formed between replicated daughter DNA molecules ^4^. These intertwines need to be removed before replicated chromosomes can segregate during cell division ^7^. To overcome the topological limitations associated with the DNA double helical structure cells have evolved a class of enzymes, DNA topoisomerases, which alleviate torsional strain by introducing transient breaks on the DNA ^8^.

There are two main types of topoisomerases, termed type I and type II, depending on the number of DNA strands they cleave and whether they require ATP ^9^. Unlike type I topoisomerases that cleave only one DNA strand, type II topoisomerases are characterized by their ability to cleave both DNA strands and use the energy derived from ATP hydrolysis in their reaction. Both type I and type II can alleviate the build-up of torsional strain due to supercoiling, but only type II can resolve intertwining between two DNA molecules ^9^.

Type II topoisomerases are functional dimers capable of passing one segment of DNA (the T-segment) through a transient double-stranded break that they make (by trans-esterification reactions where the 5′ ends of the strands are covalently linked with a pair of conserved tyrosine residues) in a second segment of DNA (the G-segment) using a gated mechanism ^10^.

Human cells contain two isoforms of the type II topoisomerase (TOP2), TOP2α and TOP2β, ^11^ differing in the polypeptide sequence of their carboxy-terminal domains (CTDs). TOP2α has been associated with DNA replication and mitosis, ^12^ whereas TOP2β has been linked to transcriptional regulation of gene expression in differentiated cells ^13,14^.

TOP2α is cell-cycle regulated, increasing in activity from mid-S phase until mitosis and decreasing rapidly upon mitotic completion ^15^ and thus thought to be the main enzyme removing DNA intertwines in mitosis. It acts along the chromosome arms prior to the onset of metaphase ^1,2^ and at the centromere at the anaphase onset ^3^. TOP2β is enriched at boundaries of topologically associated domains (TADs) ^16^ suggesting that it resolves topological stress arising during genome folding in interphase nuclei.

Before the discovery that cohesin complexes are responsible for sister chromatid cohesion ^17,18^, intertwines were thought to be the main means of sister chromatid cohesion, maintaining the physical proximity of chromatids until mitosis ^19^. Interestingly, cohesin-bound sites are also thought to contain sister chromatid intertwines (SCIs) ^20,21^, however, it is unclear whether the presence of cohesin at these sites promotes concatenation or inhibits decatenation of SCIs by TOP2 during interphase. Consistent with this view, TOP2α-dependent decatenation of centromeric regions in Hela cells occurs only after cohesin is removed by separase at the onset of anaphase ^3^.

Many studies have investigated the roles of TOP2α and TOP2β using bulk biochemistry, but these studies used heterogeneous DNA substrates, potentially obscuring detailed enzymatic mechanisms. In this study, we have used correlative single-molecule quadruple-trap optical tweezers with fluorescence detection ^22,23^ to generate braided pairs of DNA molecules and have studied their resolution by human TOP2α. Our data show that TOP2α resolves single and multiple braids rapidly and efficiently when loaded near to, or at the junction between the two interlocked DNAs, but is unable to do so when pre-loaded on linear DNA with ATP. Lastly, we show that cohesin binding to DNA braids regulates TOP2α activity by blocking substrate accessibility and impeding resolution.

## Results

### Generating DNA braids to study TOP2α activity in the optical tweezers

To study the activity of human TOP2α resolving intertwines, we used a correlative quadruple-trap optical tweezers with confocal fluorescence detection (Q-Trap)^22,23^. The instrument has four optical traps allowing the capture of two independent dsDNA molecules and their manipulation in three dimensions. We conducted our experiments in a multi-channel laminar flow cell, where the beads and captured DNAs can be moved across different channels containing distinct protein complexes and buffers (Fig. 1a). We used up to six defined channels of the flow cell. Channel 1 and 2 contained streptavidin-coated polystyrene beads and λ-DNA molecules (48.5 kb) with biotinylated ends respectively, whereas channels 3 to 6 contained different buffers and proteins depending on the experiment (Fig. 1a).

**Figure 1:**
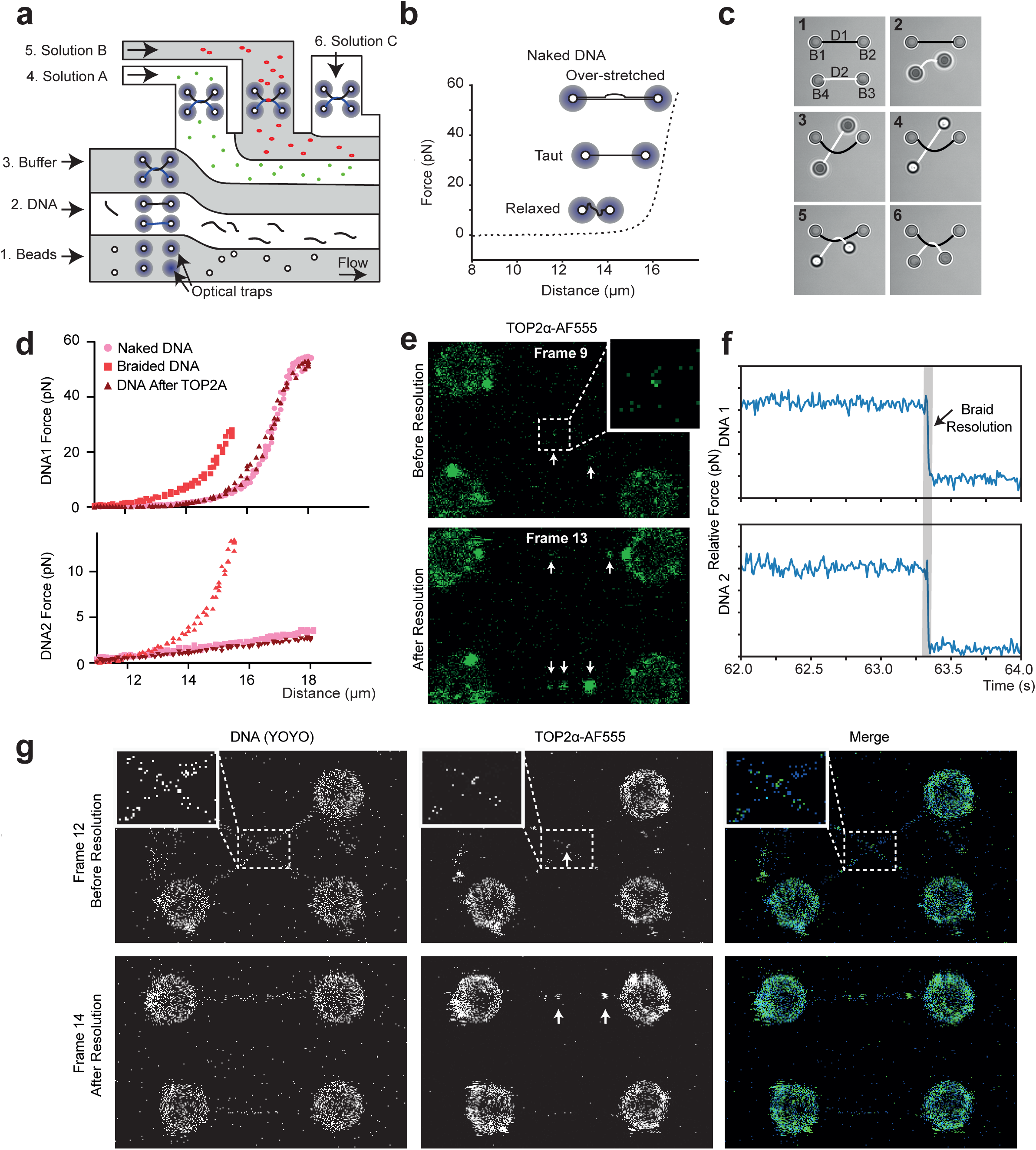
Single molecule visualisation of TOP2α DNA braid resolution. **a,** Schematic of microfluidic flow cell used for single molecule assays. Four beads can be captured by optical traps in channel 1, a DNA can be caught between pairs on beads in channel 2, force extension curve collection and DNA braid is performed channel 3 and different experimental solutions can be added to channel 4, 5 and 6. **b,** Example of force extension curve of naked DNA following Worm Like Chain model, indicating DNA conformation at different forces. **c,** Images from bead movie (Sup. Movie 1) with drawn DNA illustrating how a DNA braid is produced. DNA1 and 2 are held between beads 1 and 2, and 3 and 4, respectively. Bead 3 and 4 are moved up in Z, bead 3 is passed over DNA1, then moved down in Z before being passed under DNA1. **d,** Force extension curves in the absence of YOYO, pulling on DNA 1, measuring force experienced by DNA1 or 2. Naked DNA1 has expected contour length of ∼16μm, while pulling on DNA1 results in no force in DNA2. After braiding, DNA1 contour length is shorter and pulling on DNA1 induces force in DNA2. After TOP2α treatment, DNA returns to that of naked DNA. **e,** Direct visualisation of braid resolution by TOP2α-AF555, before resolution (frame 8) TOP2α-AF555 is in centre of 4 beads, and after resolution (frame 13), TOP2α-AF555 between bead 1-2 and 3-4. **f,** Force vs time curve of the braid resolution event shown in **e,**. **g,** Direct visualisation of braid resolution by TOP2α-AF555 in the presence of YOYO, before resolution (frame 12) braid can be observed as cross using YOYO, with TOP2α-AF555 at the junction and after resolution (frame 14) parallel DNA with bound TOP2α-AF555 can be observed. White arrows indicate bound TOP2α-AF555, scale is provided by 4.3 μm beads.

To confirm capture of individual λ-DNA molecules between each bead pair, the distance and force between the bead pair is recorded starting with the beads in proximity and increasing their relative distance. This generates force-extension (FE) curves (Fig. 1b). When a DNA is captured between the bead pair, the FE curve can be described by the Worm-Like Chain model, where a sharp increase in force is observed when the contour length of the DNA is reached (Fig. 1b). For λ-DNA molecules (48.5 kb) this is observed when beads reach a distance of ∼16μm (Fig. 1b).

To generate braided DNA substrates, two pairs of beads (B1/B2 and B3/B4, Fig. 1c) are initially trapped in channel 1 using the four optical tweezers, and then transferred to channel 2, where two DNA molecules (D1 and D2) are captured between the respective bead pairs (Fig. 1c and Movie 1). The two DNA molecules are then moved to channel 3 containing buffer (Fig. 1a), where FE curves are measured to confirm the presence of intact individual DNA molecules between each bead pair. To generate a braid between D1 and D2, we first place the two DNA molecules parallel to each other on the same optical Z-plane (Fig. 1c, frame 1). D2 is then moved below the optical plane and B3 with D2 attached, is passed under D1 (Fig. 1c, frames 2-3). At this point, D2 is raised above the optical Z-plane (Fig. 1c, frame 4), B3 is moved over D1 (Fig. 1c, frame 5), and finally, B3 and B4 are lowered back into the optical Z-plane, such that both D1 and D2 are in the focal imaging plane (Fig. 1c, frame 6). This sequence of manipulations results in a single braid between the two captured DNA molecules that contains a right-handed crossing (Fig. 1c). Since D1 and D2 are torsionally relaxed and free to rotate because they are attached to the streptavidin beads through biotin on one of the DNA strands, this protocol produces braids with similar characteristics to CatA-type catenanes ^24^.

Braids can be detected by monitoring changes in FE curves before and after their generation (Fig. 1d). Before braiding, FE curves between B1 and B2 show the expected Worm-Like Chain profile, characteristic of λ-DNA molecules (Fig. 1d; Naked DNA). However, the generation of a single braid results in a shorter DNA contour length (Fig. 1d; Braided DNA), as expected from the coupling of the two DNA molecules.

### TOP2**α** efficiently resolves DNA braids

Next, we sought to investigate whether human TOP2α was able to resolve DNA braids generated on the quadruple-trap. To this aim, we purified active human TOP2α and labelled it with Alexa Fluor 555 (AF555) (Sup Fig. 1a-c). Following the formation of a single braid in channel 3 as described (Fig. 1c), we moved the DNAs into channel 4 containing 2 nM TOP2α-AF555 and 1 mM ATP. We briefly flowed fresh protein through the channel and then incubated the DNAs with protein. TOP2α-AF555 often appeared briefly at the middle of the 4 beads, where the braid was expected (Fig. 1e; frame 9, Sup Fig. 2a; Before resolution), and rapidly moved either between B1-2 or B3-4 (Fig. 1e; frame 13, Sup Fig. 2a; After resolution, Sup. Movie 2). The Worm-Like Chain profile, characteristic of single λ-DNA molecules between each bead pair, was detected after braid resolution (Fig. 1d; DNA after TOP2α, Sup Fig. 2b). Moreover, force during the experiment confirmed the resolution of the braid. The force exerted on D1 and D2 exhibited simultaneous sharp drops at the moment when resolution takes place (Fig. 1f, Sup Fig. 2c), as expected when uncoupling the two DNA molecules.

**Figure 2:**
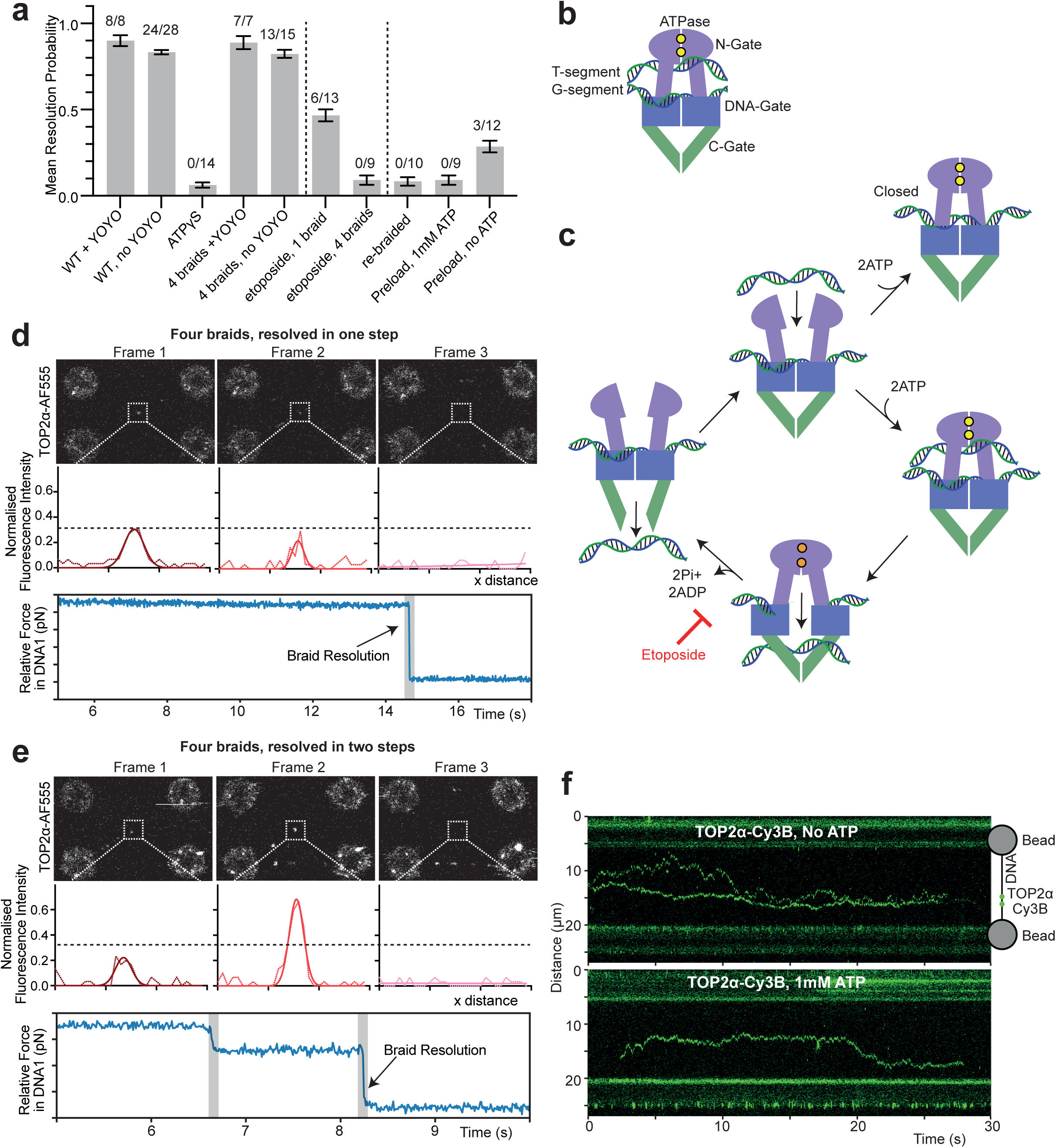
Interrogating the TOP2α enzymatic cycle. **a,** Mean proportion of DNA braid resolution by TOP2α-AF555 in different conditions. Resolution events out of total sample size indicated in each case, error bars indicate one standard error. **b,** Schematic of TOP2α enzyme with two DNA segments. **c,** Diagram showing current understanding of TOP2α enzymatic mechanism. **d, e,** Force vs time curve of 4 braid resolution, occurring in one or two steps, respectively, with corresponding scan images and TOP2α-AF555 intensity. **f,** Kymographs showing TOP2α-Cy3B associating and diffusing on DNA in the absence (top) and presence (bottom) of ATP. Scale is provided by 4.3 μm beads.

We also followed TOP2α-mediated resolution of the two DNAs by imaging DNAs in the presence of YOYO-1 dye (Fig. 1g, Sup Fig. 2a) ^25^. Holding the DNA molecules at ∼5 pN, we could visualize the braid between D1 and D2, which appeared as a cross near the middle of the 4 traps (Fig. 1g, frame 12 and Sup Fig. 2d; before decatenation). This structure was rapidly resolved into two fully separated DNA molecules (Fig. 1g; Frame 14, Sup Fig. 2d, Sup. Movie 2) consistent with resolution by TOP2α. Importantly, in many cases, we were able to visualize the binding of TOP2α-AF555 to the DNA junction just ahead of its resolution (Fig. 1g, Sup Fig. 2d). We thus conclude that TOP2α efficiently resolves a single DNA braid generated between the pair of λ-DNA molecules in our experimental setup.

### TOP2**α** resolution of DNA braids requires ATP hydrolysis

Our assay demonstrated robust braid resolution by TOP2α (mean resolution probability of 0.90 ± 0.03 and 0.83 ± 0.01, with and without YOYO, respectively, Fig. 2a). However, type II topoisomerases like TOP2α require ATP for activity ^26^ (Fig. 2b-c). The generally accepted model is that ATP binding closes the N-terminal gate and is sufficient for transport of the T segment through the G segment ^27–29^. In contrast, ATP hydrolysis is thought to induce reopening of the N-terminal gate closing the cycle ^30^. It is however unclear, whether ATP binding is sufficient for release of the transported T segment through the C-terminal gate. We sought to test whether DNA braid resolution requires ATP hydrolysis. To test this, we used ATPγS, a non-hydrolyzable ATP analogue. Although we observed TOP2α localisation at the DNA junction, no resolution events were observed when ATPγS was present (N=14, Fig. 2a, Sup Fig. 3a). Therefore, we confirm that resolution in our assay requires ATP hydrolysis by TOP2α and conclude that while ATP binding might be sufficient for transfer of the T-segment through the broken G segment ^27–29^, release of the DNA through the C-terminal gate does not occur in the absence of ATP hydrolysis.

**Figure 3:**
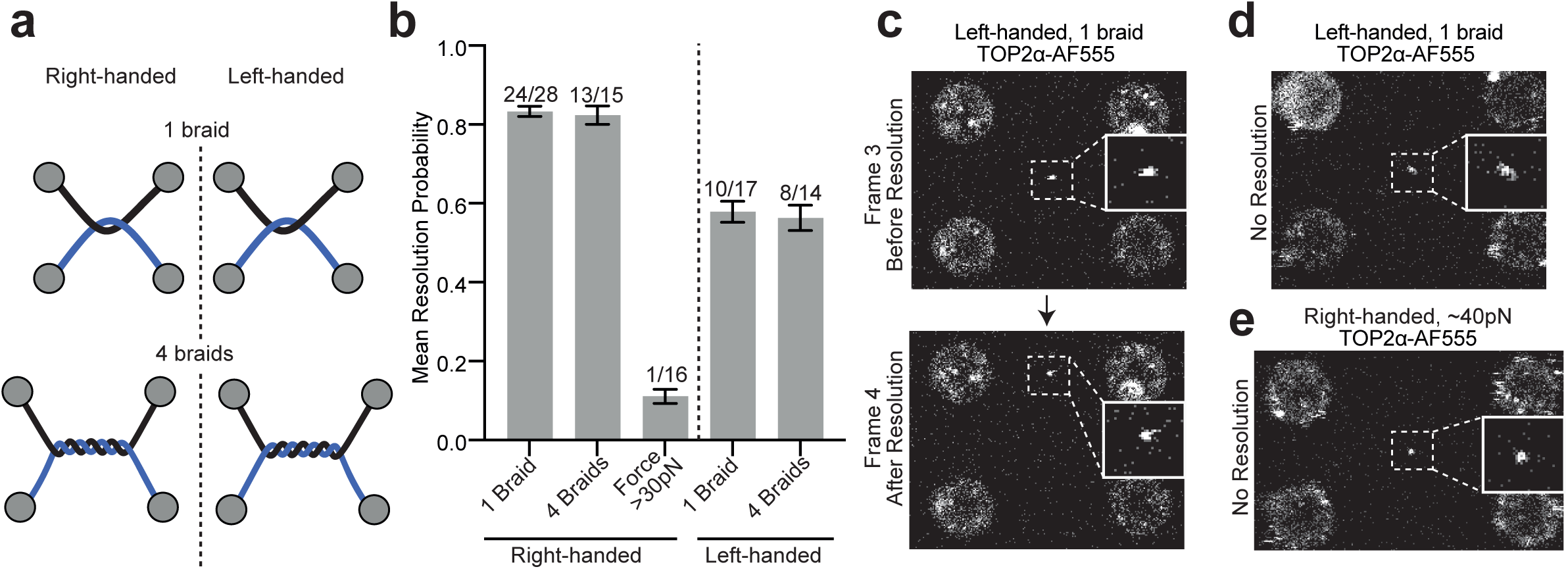
TOP2α DNA braid resolution is affected by DNA conformation. **a,** Schematic of right- and left-handed DNA braids. **b,** Mean resolution probability by TOP2α-AF555 in the absence of YOYO-1of left- and right-handed substrate, and right-handed substrate held at high forces of 30-50 pN. Resolution events out of total sample size indicated in each case, error bars indicate one standard error. **c,** Example of left-handed braid resolution event, before resolution (frame 3) TOP2α-AF555 is in centre of 4 beads, and after resolution (frame 4), TOP2α-AF555 between bead 1-2. **d,** Example of left-handed braid that did not resolve. **e,** Example of right-handed braid held at 40 pN that did not resolve.

### TOP2α resolves DNAs with multiple braids

Previous work has suggested that individual TOP2α dimers can deal with multiple DNA crosses in supercoiled templates in a processive manner ^31^. In our assays, TOP2α efficiently removes a single braid between DNAs (Fig.1 d-g). We next tested whether intertwining with higher numbers of braids could also be resolved efficiently in this assay. By repeating the DNA braiding procedure, we made DNA substrates with up to four right-handed braids in the buffer channel. Then, we incubated the braided DNAs in channel 4 with TOP2α and ATP, with and without YOYO-1(Fig. 2a, Sup Fig. 3b). We observed that the efficiency of resolution of DNAs with multiple braids is like that of DNAs containing a single braid (Fig. 2a). Interestingly, we were able to detect either one or two drops in force, when monitoring the traps during the resolution of multiple braids (Fig. 2d-e). When resolution exhibited one drop in force, in most cases, there was a consistent fluorescence intensity at the junction (Fig. 2d). However, in three cases, we detected two drops in force which was coupled with an increase in TOP2α fluorescent intensities over time (Fig. 2e), suggesting that more than one enzyme was involved in these cases. These results suggest that while multiple TOP2α can work together, a single bound complex can also act in a processive manner when resolving multiple braids in proximity, just as it has been demonstrated for supercoil relaxation ^31^.

### Etoposide inhibits TOP2α activity

TOP2α is the target of the chemotherapy drug etoposide. Previous research has established that etoposide impedes decatenation through its ability to bind to and stabilize the DNA cleavage intermediate (Fig. 2c). Next, we sought to test TOP2α activity in our assay in the presence of etoposide. First, we investigated the effect of etoposide on TOP2α resolution of substrates containing a single DNA braid. In the presence of 50 μM etoposide, the mean probability of resolution dropped from 0.83 ± 0.01 to 0.47 ± 0.01 (Fig. 2a). This result is consistent with a mechanism by which etoposide stabilises the cleavage intermediate and prevents release of the T-DNA segment through the C-terminal gate, thereby blocking resolution. We reasoned that increasing the number of DNA braids between the two DNAs should reduce the resolution probability further, as it gives the etoposide multiple catalytic cycles to bind in. To investigate this, we introduced four braids between the DNAs and tested TOP2α resolution of these interlocked DNAs in the presence of etoposide. We never observed resolution of these braided molecules (N = 0/9, Fig. 2a). We conclude that, as expected, etoposide inhibits TOP2α activity resolving braided DNAs in our single-molecule assay.

### TOP2α DNA braid resolution requires controlled loading to the DNA junction

In our experiments, we observed that TOP2α remained bound to the DNA after the resolution events (Fig. 1e and 1g; frame 14). Since TOP2α was able to resolve DNAs with multiple braids efficiently (Fig. 2a), we decided to test whether TOP2α molecules that stayed bound after a resolution event could mediate a second round of resolution when a new braid is introduced between the same DNA molecules. To this aim, a new braid was generated as described (Fig. 1c) after an initial round of resolution by TOP2α. Surprisingly, we did not observe the unlinking of these re-braided molecules despite the continued presence of TOP2α on the DNAs and ATP, including near the braid itself (Fig. 2a, Sup Fig. 3c). This data demonstrates that following a round of resolution, bound TOP2α cannot untangle a newly introduced DNA braid. This is surprising since our experiments with DNAs containing multiple braids demonstrate the enzyme is capable of processive resolution of more than one junction (Fig. 2c-d). These results suggest that loading at the DNA braid might be important for efficient TOP2α resolution.

To test this further, we decided to separate the loading and braiding steps. First, we confirmed that TOP2α can load onto dsDNA molecules (Fig. 2f) by incubating a single piece of dsDNA in a channel with TOP2α-Cy3B. Kymograph analysis shows that TOP2α molecules not only bind but also freely diffuse along dsDNA. Importantly, this behaviour was observed both in the presence and absence of ATP (Fig. 2f). Next, we investigated whether these TOP2α molecules loaded to linear D1 and D2 were able to resolve a newly generated DNA braid. To this aim, we incubated D1 and D2 before braid formation at the junction of channels 3 and 4, containing 2 nM TOP2α-AF555 and 1 mM ATP. After observing that TOP2α bound the individual DNAs efficiently, we moved D1 and D2 with bound TOP2α, and generated a single braid in channel 3 before incubating in channel 5 with 1 mM ATP and YOYO-1 dye. Pre-loading of TOP2α in the presence of ATP completely prevented resolution events, despite TOP2α localising at the junction (Fig. 2a, Sup Fig. 3d-e). This is consistent with the observation that TOP2α could not resolve newly formed braids after a round of resolution had taken place (Fig. 2a). This result suggests that TOP2α needs to load at, or very near, the braid between the two DNA molecules with ATP to resolve it, and that enzyme preloaded on linear dsDNA with ATP is unlikely to remove braids, even if these are encountered during linear diffusion.

Since TOP2α was able to bind linear DNA in the absence of ATP (Fig. 2f), we decided to test whether the absence of ATP impacted on the ability of pre-loaded TOP2α to resolve braids. To this aim, we pre-loaded TOP2α omitting ATP in the buffer and then generated a single braid before moving DNAs to channel 5 with 1 mM ATP and YOYO. We observed that under this regime TOP2α had a mean resolution probability of 0.29 ± 0.03 (N = 3/12, Fig. 2a, Sup Fig. 3f). These results suggest that TOP2α molecules bound to dsDNA maybe less able to capture the second DNA necessary to proceed with resolution of the braids. This might be due to the fact that pre-bound TOP2α sits on dsDNA with a geometry that is incompatible with the capture of the second DNA, or alternatively, that ATP binding and closure of the N-terminal gate, when only one DNA molecule has been captured, generates a non-productive enzyme-DNA complex that is both unable to capture a second DNA (with a closed N-terminal gate, Fig 2c) or proceed to reopen the N-terminal gate. This hypothesis might explain why loading in the absence of ATP allowed a small percentage of the newly generated DNA braids to resolve (Fig. 2a).

### TOP2**α** resolves right-handed crosses more efficiently than left-handed

In our experiments, depending on the direction that the DNAs are braided they can contain a left-handed or a right-handed cross (Fig. 3a). When generating the braids, the movement of bead 3 dictates the handedness of the cross. When B3 is first moved under D1, the cross is right-handed, while when B3 is first moved over D1 the cross is left-handed (Fig. 3a). Previous studies have shown that human TOP2α removes multiple DNA braids present on the same molecule at a faster rate when they contain right-handed crosses ^32^. However, this was only observed when the DNA braids were held at a very low tension, <0.4 pN ^32^. In the current configuration, our instrument does not allow us to perform resolution experiments at low tensions. However, it allows us to see the outcome of every TOP2α that binds, hence we sought to compare the relative efficiency of TOP2α resolving DNA molecules linked with a single right or left-hand braid held at moderate tensions (∼5 pN). TOP2α removed DNA braids crossed in both directions, however, the efficiency was higher for right-hand crosses (Fig. 3b-d). Next, we extended our analysis to DNAs with multiple braids. We compared the efficiency of TOP2α resolution for DNA molecules containing four right-handed or four left-handed braids. As observed for a single DNA braid, TOP2α exhibited higher efficiency on DNAs containing multiple braids with right-handed crosses (Fig. 3b). These results demonstrate that in our assay TOP2α has a chiral preference for right-handed crosses, as previously reported ^32^.

### TOP2α DNA braid resolution is inhibited with force

Previous studies showed the unlinking rate was dramatically reduced with tension ^32^. In our assays, efficient resolution was observed when the DNA braids were subjected to a mean force of ∼5 pN (Fig. 3b). Mitotic chromosomes are pulled apart by kinetochore attachments, each of which having a mean rupture force ∼10 pN ^33^. Since each kinetochore contains 15-25 microtubule attachments, the centromeric intertwines can be subjected to 100s of pN of force. Given this, we sought to investigate the upper force limit for efficient TOP2α resolution of DNA braids. After introducing a single right-handed DNA braid between a pair of DNA molecules, we pulled on beads 1 and 2, generating an average force at the DNA braid between 30-50 pN, prior to moving it to channel 4 with TOP2α and ATP. Resolution was significantly inhibited at this force range, resulting in a mean resolution probability of 0.11 ± 0.02 (1 out N=16, Fig. 3b, e), compared to 0.83 ± 0.01 for ∼5 pN. These results confirm that DNA braid resolution decreases with increased tension, as previously demonstrated ^32^, and suggests that TOP2α is functional at a range of forces that are consistent with those expected to occur on segregating mitotic chromosomes.

### Cohesin inhibits TOP2α DNA braid resolution

Despite the advances in understanding the molecular mechanisms behind the enzymatic activity of TOP2α, little is known about the regulation of accessibility to DNA substrates by other proteins. Cohesin is a multi-subunit complex that organizes interphase chromatin and provides sister chromatid cohesion ^34^. Cohesin forms a tripartite ring structure with compartments that can trap one or two DNA molecules ^35–37^. Precisely how cohesin does this is unknown. Interestingly, binding of yeast cohesin to chromosomes correlates with the presence of intertwines between sister chromatids ^20,21^. Moreover, the resolution of catenation by TOP2α has been shown to follow cohesin removal from centromeric regions on mammalian chromosomes ^3,38^, raising the possibility that cohesin prevents TOP2α from accessing intertwines. With this in mind, we sought to investigate whether the presence of cohesin in our assays would affect the resolution of the DNA braids by TOP2α.

First, we purified human cohesin (SMC1, SMC3, RAD21 and STAG1-ybbr) and MBP-ΔN-NIPBL and confirmed activity with ATP hydrolysis assays (Sup. Fig. 1a-b, d). We used the ybbr tag to label the complex with ATTO647N(A647N). We then investigated how cohesin interacts with braided DNA. We generated a right-handed braid in channel 3 and moved the structure to channel 4 containing 2nM cohesin-A647N, 4 nM MBP-ΔN-NIPBL, 1 mM ATP and 0.1 nM YOYO. Cohesin complexes were found to load onto both dsDNA regions as well as the DNA junction (Fig. 4a). When we tracked cohesin over time, kymographs showed that cohesin bound to the DNA junction (braid) remained immobile, while cohesin on dsDNA exhibited diffusion along the DNAs (Fig. 4a). After cohesin was bound to the DNA braid, we moved the DNAs to channel 5 containing TOP2α, 1 mM ATP and 0.1 nM YOYO-1 and tested whether resolution of the DNA braids took place. When pre-incubated with cohesin resolution by TOP2α was dramatically reduced to a mean resolution probability of 0.25 ± 0.03 (3 out of N=14), compared to 0.90 ± 0.03 (8 out N=8) observed for TOP2α without cohesin (Fig. 4b), despite TOP2α being bound near cohesin at the DNA junction (Fig. 4c).

**Figure 4:**
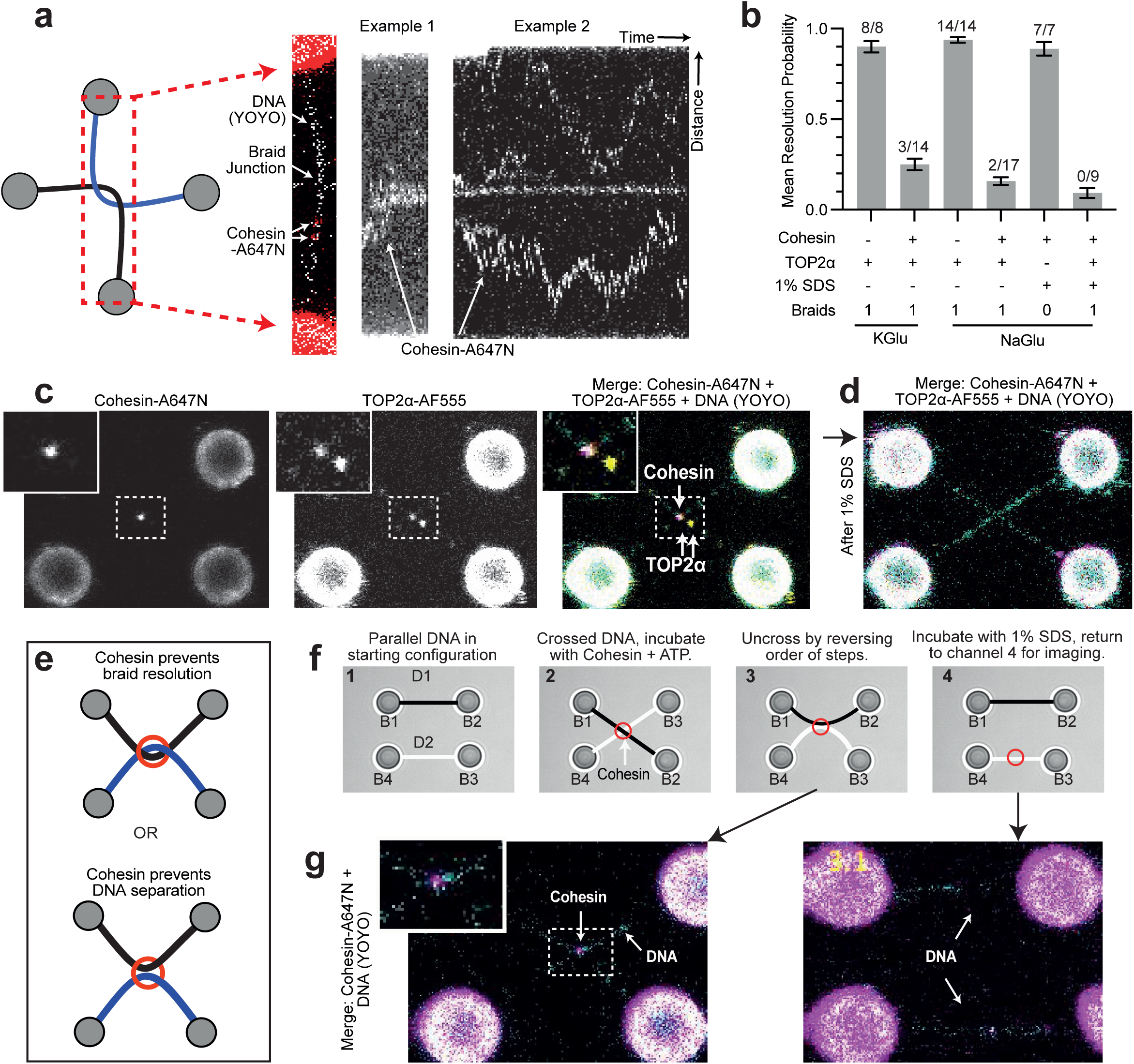
Cohesin blocks TOP2α DNA braid resolution. **a,** Cohesin-A647N /MBP-ΔN-NIPBL kymograph on braided DNA substrate. Kymograph is generated from scanning a limited area of reorientating beads. **b,** Mean proportion of DNA resolution in experiments with cohesin-A647N and TOP2α-AF555. DNA resolution events out of total sample size indicated in each case, error bars indicate one standard error. **c,** Cohesin-A647N localised at a catenation junction with TOP2α-AF555. **d,** DNA braid incubated with cohesin-A647N then TOP2α-AF555 is not resolved after incubating with 1 % SDS. **e,** Two hypothetical modes of how cohesin could prevent DNA resolution. **f,** Schematic of cohesin bridging assay. DNA1 and 2 (D1 and D2) are crossed by swapping the positions of bead 2 and 3 (B2 and B3) and incubated with cohesin-A647N/MBP-ΔN-NIPBL, 1mM ATP and 0.1nM YOYO. B2 and B3 are then returned to original positions, resulting in cohesin bridging D1 and D2. Bridging can be resolved with 1% SDS, before imaging with YOYO. **g,** Example images of cohesin-A647N DNA bridging from step 3 and 4 shown in **f,**. Scale in all scan images is provided by 4.3 μm beads.

These results suggest that cohesin binding to the DNA braid reduces resolution efficiency by TOP2α drastically. Previously work from our group and others have shown that yeast cohesin can tether two DNA molecules brought into proximity ^23,37^, raising the possibility that cohesin may prevent resolution of the two DNAs by bridging, rather than preventing DNA braid resolution by TOP2α (Fig. 4e). We found that human cohesin-A647N and MBP-delN-NIPBL are also able to tether two DNAs and that this bridging is protein mediated, as it can be efficiently dissolved by 1% SDS (resolving 7 out of N=7) (Fig. 4b, f-g, Sup. Movie 3). Therefore, to confirm that cohesin is preventing TOP2α-mediated resolution of the DNA braid, rather that blocking DNA separation by bridging the two DNAs, we included a control to dissolve cohesin tethering between the DNAs after incubation with cohesin and TOP2α. To this aim, we generated a single DNA braid in channel 3, incubated with cohesin-A647N and MBP-delN-NIPBL with 1mM ATP in channel 4 and TOP2α-AF555 with 1mM ATP in channel 5, and then moved the structure to channel 6 which contained 1% SDS. This treatment did not result in resolution of the crossed DNAs, demonstrating that the DNAs remained braided (0 out of N=9) (Fig. 4b-d, f, Sup. Movie 4). Therefore, we conclude that the presence of cohesin at the DNA cross compromises the ability of TOP2α to resolve the braid linking D1 and D2 (Fig. 4b).

## Discussion

Numerous studies have investigated the function of TOP2α and TOP2β ^39^. Defining the enzymatic details of their activity has contributed significantly to our understanding of their fundamental roles in various biological processes. However, most studies have analysed TOP2 activities using bulk biochemical analysis, which often involves DNA substrates that lack homogeneity and contain a variable number of topological features, for example a range of catenanes. This inherent heterogeneity in the substrates can potentially obscure some fine details of the enzymatic mechanisms at play.

Our assay provides multiple outputs to measure the release of T segment DNA in native conditions. Previously, type II topoisomerases have been shown to transport the T DNA through the cleaved G DNA in the absence of ATP hydrolysis ^27–29^. In such bulk assays, decatenation of singly catenated plasmids, in the presence of a non-hydrolyzed ATP analog was achieved after washes with high ionic strength buffers (1M NaCl)^27–29^. We did not observe DNA braid resolution when we substituted ATP for ATPγS in our assays (Fig. 2a), despite the fact that TOP2α was able to bind to DNA braids (Sup Fig 3a). These data suggests that while ATP hydrolysis might not be necessary for transport of the T DNA through the cleaved G DNA, it is likely to be important for the release of the T DNA through the C-terminal gate. Our assays using etoposide are consistent with this. Previous work has suggested etoposide acts before the release of the final ADP, allowing transit of the T-segment, but prevents re-ligation of DNA, resulting in decatenation of single catenated plasmids in assays where the reaction is followed by is denaturing condition ^40^. In our assays, etoposide is able to inhibit resolution of a single braid, suggesting it prevents release of T DNA from the C-gate. Consistent with this, others have reported etoposide can promote TOP2α DNA looping, potentially by preventing T DNA release ^31^. Collectively this suggests TOP2α could have a release checkpoint to ensure G DNA is re-ligated before release of T DNA, where it has been previously demonstrated that G DNA can be re-ligated while T DNA is trapped in a covalently closed C-gate ^41^.

Another advantage of our assay is that it enables the examination of TOP2α enzymatic function in substrates with a defined number of DNA braids (Fig. 1c). We have contrasted TOP2α mediated resolution of DNAs with one and four braids (Fig. 2a) and found that TOP2α molecules can resolve both with similar efficiency (Fig. 2a). Despite being able to resolve multiple braids, we found that loading of TOP2α onto a single dsDNA, particularly in the presence of ATP, compromises the ability of the enzyme to resolve newly formed DNA braids (Fig. 2a). This result raises the possibility that TOP2α acts through a carefully regulated loading mechanism to ensure both DNA segments are grasped correctly. In our assays, TOP2α pre-bound to dsDNA molecules might sit on a configuration where the geometry of the enzyme-DNA interaction prevents the second DNA from entering through the N-terminal gate as the enzyme encounters it during diffusion on the bound DNA molecule. The observation that TOP2α pre-bound to dsDNA can in the absence of ATP resolve newly formed DNA braids, although with reduced efficiency (Fig. 2a), raises the interesting possibility that ATP binding by TOP2α molecules that have captured a single DNA might generate an intermediate where ATP-dependent closure of the N-terminal gate blocks capture of the second DNA generating a situation where the enzyme stalls at a non-productive intermediate (closed state, Fig. 2c). We envisage that in substrates with multiple DNA braids in proximity, at each resolution event the enzyme has a high probability to capture two DNAs simultaneously for the next enzymatic cycle, before ATP closure of the N-terminal gate, to proceed efficiently, thus explaining the processivity observed (Fig. 2a).

Our observations not only provide several important insights into TOP2α action but also open up important questions of its regulation in cells. Firstly, it was previously unclear how TOP2α accessed the DNA crosses it resolves, whether it loaded onto dsDNA and found the DNA junctions by moving along the DNA, or whether it needed to load at the DNA cross. Our results indicated that TOP2α activity is more efficient when the enzyme loads at or near the junctions, since this maximises the probability of capturing both DNAs. Moreover, we observed that TOP2α remained stably associated both in the presence and absence of ATP when loaded onto dsDNA (Fig. 2f) but was unable to resolve newly formed DNA braids in the presence of ATP (Fig. 2a), suggesting that the enzyme might get stuck on dsDNA in a non-productive configuration. These findings have important regulatory implications related to substrate accessibility *in vivo.* Previous studies suggest chromatin reduces TOP2α accessibility ^42^, and that direct interaction between TOP2α the BAF chromatin remodeller allows recruitment ^43^. Alternatively, these findings could point to non-catalytic, structural roles for TOP2α, where cell-based studies have demonstrated that TOP2α degradation results in a chromosome decompaction phenotype which is not observed using simple TOP2α inhibition ^44^.

Another interesting observation from our study is the chiral preference to resolve right-handed DNA braids exhibited by TOP2α (this study and ^32^). Intertwining between sister chromatids is a consequence of the late stages of DNA replication ^4^. Due to the limited space between converging forks, topoisomerase activity in front of the replication forks is compromised and fork swivelling is thought to be the preferred method to relieve superhelical tension at replication termination sites ^45^. The consequence of fork rotation is the formation of DNA crosses between replicated sister chromatids behind the forks, or precatenanes; these become sister chromatid intertwines (SCIs), or full catenanes, upon replication completion. Importantly, because of the right-handed helicity of DNA ^5^ fork swivelling produces right-handed crosses behind the fork and consequently sister chromatid intertwines are wound around each other in a right-handed manner. Since TOP2α is likely to be the primary mitotic decatenase ^1–3^ the chiral preference for right-handed crossing might enables the enzyme to be more effective at removing catenanes formed during DNA replication.

Our results demonstrate that loading to, or close to, DNA crosses is a key determinant for rapid and successful decatenation by TOP2α. Our finding that cohesin complexes stably associate with braided junctions and have an inhibitory effect on TOP2α resolution of DNA braids (Fig. 4a,b) raises important insights into genome segregation. Interestingly, the presence of SCI in yeast chromosomes requires cohesin ^20,21^ and it has been well characterised that human centromeres are highly catenated regions whose decatenation by TOP2α occurs during anaphase ^46^, only after separase dependent removal of cohesin ^3^. Collectively, these observations indicate that the presence of cohesin on catenated regions of the genome might prevent SCI resolution by topoisomerase II. We propose that cohesin inhibition of TOP2α could occur by restricting simultaneous capture of the two DNAs at catenation crosses, which is an important requirement for efficient resolution of DNA braids by TOP2α.

In summary, we provide valuable insights into the possible regulation of TOP2α recognition of catenated substrates. We propose that rather than a mere passive presence within the nuclear milieu, randomly encountering and resolving DNA crossings, TOP2α function is likely to rely on a highly regulated orchestration of the enzyme access to specific DNA substrates. We anticipate that this regulation will include mechanisms that restrict access to regions where processing should be avoided, such as cohesion sites, as well as regulation that facilitates the presentation or capture of DNA crosses. Given the importance of human TOP2 enzymes as therapeutic targets and their increasingly recognized role as a potential source of genome instability, understanding how their activities are controlled and restricted to proper sites of action is an important question for the future.

## Contributions

E.C. and L.A. designed experiments. E.C. cloned, expressed, purified all proteins. E.C. performed experiments and analysed the data. S.S. performed experiment with multiple left-handed braids. G.F. designed cohesin bridging assay. D.R. designed chirality assays. L.A. wrote the first draft. L.A., D.R. and E.C. reviewed the manuscript. All authors contributed to editing and provided additional text for the manuscript.

## Acknowledgments

We thank all members of the DNA motors group for discussions, comments on the manuscript and generation of communal reagents. We thank all members of the Single molecule imaging group for discussions.

## Methods

### Protein purification

Human TOP2α was encoded from the pLIB vector with a C-terminal 3C-ybbr-tev-strepII tag, Cohesin STAG1 tetramer from pBIG2ab ^47^ with a C-terminal 3C-His10 tag on SMC3 and a C-terminal 3C-ybbr-tev-strepII on STAG1 and NIBPL with a deletion of N-terminal 1162, an N-terminal MBP and C-terminal 3C-ybbr-tev-strepII tag from pLIB. All constructs were transposed into DH10EMBacY and purified Bacmid transfected into SF9 cells. After 72 hours, virus was harvested and further amplified in SF9 cells before being used for expression in either SF9 or HighFive cells for 72 hours. Cell pellets were resuspended in purification buffer (20 mM HEPES [pH 8], 300 mM KCl, 5 mM MgCl2, 1 mM DTT, 10% glycerol) supplemented with 1 Pierce protease inhibitor EDTA–free tablet (Thermo Scientific) per 50mL and 25 U/mL of Benzonase (Sigma) and lysed with a dounce homogeniser followed by brief sonication. Lysate was cleared with centrifugation before being loaded on to a StrepTrap HP (GE), washed with purification buffer and eluted with purification buffer supplemented with 5 mM Desthiobiotin (Sigma). Protein containing fractions were pooled, diluted 2-fold with Buffer A (20 mM HEPES [pH 8], 5 mM MgCl2, 5% glycerol, 1 mM DTT), loaded on to HiTrap Heparin HP column (GE), washed with Buffer A with 250 mM NaCl, then eluted with a gradient up to 2M NaCl. Finally, size exclusion chromatography was performed using purification buffer and a Superose 6 16/70 or increase 10/300 column.

Proteins were labelled using SFP transferase to attach HPLC purified dye conjugated CoA to the protein encoded ybbr tag ^48^. TOP2α was labelled with Alexa555 or Cy3B while Cohesin labelled with ATTO647N. After protein labelling, complex was purified with size exclusion chromatography using a superose 6 increase 10/300 column. Labelling efficiency of TOP2α Alexa555 and cohesin STAG1-ATTO647N were ∼30%, and improved to ∼70% for TOP2α by enriching using the Strep tag after labelling prior to gel filtration.

### Mass Photometry

The molecular mass of recombinant complexes was confirmed with a Refeyn TwoMP mass photometer. Sample was applied to a CultureWellTM gasket (GBL103250, Sigma-Aldrich) attached to a sample carrier slide (Refeyn). All samples were measured in 50 mM Tris pH 7.5, 150 mM NaCl, 2.5 mM MgCl_2_ buffer using a field of view 512 x 138 pixels, collecting 6000 frames with a collection time of 60s. The focal position and imaging conditions were set using a 12 μL buffer droplet and data was collected by adding 2 μL of sample, resulting in a final protein concentration of ∼5 nM. All data were acquired with using the Refeyn AcquireMP software and analysed using the Refeyn DiscoverMP software. Masses were calibrated using the NativeMarkTM unstained protein standard (LC0725, Thermo Scientific) to generate a calibration curve.

### Decatenation Assays

Catenated kinetoplast DNA (kDNA, Inspiralis, K1002) (200ng) was incubated with 80 nM of labelled or unlabelled TOP2α in 50 mM Tris pH 7.5, 125 mM potassium glutamate, 2.5 mM MgCl_2_, 0.5 mg/mL BSA in the presence or absence of 1 mM ATP for 30min at 37 degrees. Reaction was terminated with 3 μL of stop buffer (5% sarkosyl, 0.025% bromophenol blue, 50% glycerol) and incubated with 1.6 units of Proteinase K (NEB, P8107S) for 15 minutes at 37 degrees before resolving on 1% agarose TAE gel stained with SybrSafe (Invitrogen, S33102). Decatenated products confirmed by comparison to control decatenated and linearised kDNA standards (Inspiralis, KD100 and KL100, respectively).

### ATPase assays

ATPase assays were performed with complexes of wild type or ATPase hydrolysis deficient Q-loop mutants of cohesin with 50bp dsDNA, and with or without MBP-ΔN-NIPBL. Assays were performed using the EnzChek Phosphate Assay Kit (Invitrogen) modified for a 96 well plate format ^49^. Reactions contained 30 nM protein with 600 nM DNA. Final conditions included 1 mM ATP and a total salt concentration of 50 mM. Protein/DNA was preincubated in reaction mix without ATP for 15 min at room temperature before the reaction was started by addition of ATP immediately prior to putting it in the plate reader to track phosphate release. ATPase rate was determined using standard phosphate curve using linear fit of data in linear region.

### Single molecule braid resolution assays

Lambda DNA was biotinylation by end filling with Klenow DNA polymerase as previously described ^50^.Optical tweezer assays were carried out on a Lumicks Q-trap system, with integrated microfluids and confocal fluorescence microscopy. Flow cell was cleaned with 5% bleach, water, 25 mM Thiosulfate, water, before being blocked with 0.5% pluronic and 2mg/mL BSA in experimental buffer (50 mM Tris pH 7.5, 125 mM potassium glutamate, 2.5 mM MgCl_2_).

All assays were performed in experimental buffer with 0.5 mg/mL BSA except assays including 1% SDS, for which 125 mM potassium glutamate was substituted with 125 mM sodium glutamate. In all assays, beads were in channel 1, biotinylated lambda DNA in channel 2 and buffer in channel 3. Protein solutions of TOP2α or cohesin were used in channel 4 and 5 using a standard flow cell, and 1% SDS in channel 6 of 9 channel flow cell. TOP2α was used at 1nM for non-catenated binding kymographs and at 2 nM for all decatenation assays, and cohesin was used at 2 nM, with 4 nM NIPBL, with 1 mM ATP or ATPγS and 0.1 nM YOYO-1dye.

Trapping laser was set at 100%, splitting ∼67% across traps 1 and 2, to achieve equal trap stiffness of ∼ 0.30 pN/nm. All experiments were started by trapping one bead per trap, tethering DNA between bead pairs 1 and 2, and 3 and 4 and collecting force distance data for each DNA. DNA knots/braids were created using the Bluelake DNA knotting script written by Aafke van den Berg from Harbor Lumick (https://harbor.lumicks.com/single-script/e03478be-fa72-495f-8391-785e7a15e6e9) in combination with manual control to prevent bead collision and overstretching of DNA. The script moved the Z-height of bead 3 and 4 from focus at 2.8 μm to 1.2 μm, passing bead 3 under DNA1 held by bead 1 and 2, then to 4.4 μm, before passing bead 3 over DNA1 and returning to focus. This results in a right-handed twist, similar to that found in negative supercoiled DNA or produced by turning beads anti-clockwise. The script could also be looped multiple times, and in the reverse order, resulting in the opposite handedness.

In standard assays, beads were moved to a mean force of ∼5pN in the buffer channel prior to moving to protein channels. In higher force experiments, beads were moved to forces ∼30-50pN in buffer channel before moving to protein channel. High accuracy was not guaranteed for forces in these assays, as movement in flow cell can result in addition force and force offsets, and flow induced additional force. Protein was flowed into the channel at low flow rates either immediately before or after moving to the channel, to ensure consistent level of protein in solution.

### Data analysis

Images were acquired with 1% green laser, and 2% red and blue laser. Data was analysed in Lakeview (LUMICKS), Fiji and python using pylake.

Statistical analysis was performed utilizing Bayesian inference, where the catenation event was modelled as a Bernoulli trial with a decatenation being defined as a success. Since the likelihood for this was a binomial distribution, the probability of decatenation could be estimated as a beta distribution, Beta (α, β) ^51^. Using the beta uniform distribution, Beta (1,1) as the conjugate prior, the parameters for the above distribution could be calculated as α=k+1 and β=n-k+1, where n is the number of trial and k is the number of resolution events ^52^. In line with this analysis, the probability of resolution would have Mean = α/(α+β), and

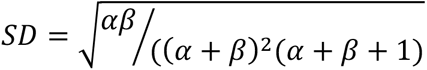

Data in figures and text is reported with mean **±** standard error.

