## Supplemental Figures 1-3 for "Substrate accessibility regulation of human TopIIa decatenation by cohesin"

**Supplementary Figure 1: Purified protein used in this study. a,** SDS page analysis of protein samples used in this study, stained either with coomassie or imaged via conjugated fluorophore. **b**, Mass-photometry analysis of proteins used in this study, indicating expected and measured molecular mass. **c,** Decatenation assay, demonstrating ATP dependent decatenation of kinetoplast DNA (kDNA) by unlabelled and labelled TOP2α. Decatenated control (Decat) and cleaved linear kDNA are loaded to demonstrate DNA is being decatenated rather than cleaved. **d,** ATP hydrolysis assay of cohesin samples in the presence of 20-fold excess of 50 bp DNA. Q indicates ATPase hydrolysis deficient mutation in the Q-loop of ATPase site. N=3, error bars indicate one standard error.

**Supplementary Figure 2: Additional examples of single-molecule visualisation of TOP2α DNA braid resolution. a,** Examples of resolution events imaged with TOP2α-AF555 in the absence of YOYO. **c, d,** Force extension curves of DNA substrate and force vs time curve of a resolution event corresponding to examples in **a,** respectively. **d,** Two additional examples of braid resolution events imaged with YOYO and TOP2α-AF555. White arrows indicated TOP2α-AF555 binding, scale in all scan images is provided by 4.3 μm beads.

**Supplementary Figure 3: Example data demonstrating TOP2α requires ATP, acts on multiple braided substrates and must be near a junction. a,** Examples of TOP2α-AF555 with 1mM ATPγS, DNA imaged with YOYO dye. Top panel shows image taken with dual excitation with blue and green laser, while bottom panel shows image taken with only green excitation to clearly illustrate TOP2α-AF555 at junction. **b,** Example data showing resolution events of substrate with four braids, imaged with YOYO and TOP2α-AF555. **c,** Example image of rebraided DNA, imaging TOP2α-AF555. **d,** Example image of preloading experiment in the presence ATP, then **e,** braiding the DNA before imaging with YOYO in the presence of ATP. **f,** Example of preloading experiment in the absence ATP (left), which is then braided (middle) before incubation with 1 mM ATP and YOYO results in braid resolution (right). Scale is provided by 4.3 μm beads.

**Supplementary Movie 1: Optical tweezer generation of braided DNA substrate.** Bright field images of four optically trapped beads being moved to create braided DNA.

**Supplementary Movie 2: Real time visualisation of DNA braid resolution by TOP2α.** Two examples of DNA braid resolution in the presence and absence of YOYO dye (cyan) by TOP2α-AF555 (yellow and white, respectively).

**Supplementary Movie 3: Cohesin form DNA bridges that are resolved by SDS.** Two crossed pieces of DNA, stained with YOYO (cyan) are incubated with cohesin-A647N (magenta) with NIPBL and ATP, until cohesin- A647N binding is observed. DNA is then uncrossed, resulting to cohesin-A647N mediated DNA bridging. The DNA bridge can then be resolved with 1% SDS, leaving two pieces of intact DNA, visible when beads are pulled apart.

**Supplementary Movie 4: Cohesin prevents TOP2α braid resolution.** Braided DNA, stained with YOYO (cyan) is incubated with cohesin-A647N (magenta) with NIPBL and ATP, resulting in cohesin-A647N localising to junction. DNA is then incubated with TOP2α-AF555 (yellow) and ATP, resulting in TOP2α-AF555 colocalising at the junction with cohesin-A647N. DNA is then incubated with 1% SDS, before returning to channel with YOYO for imaging, to reveal loss of protein from the junction and no braid resolution.

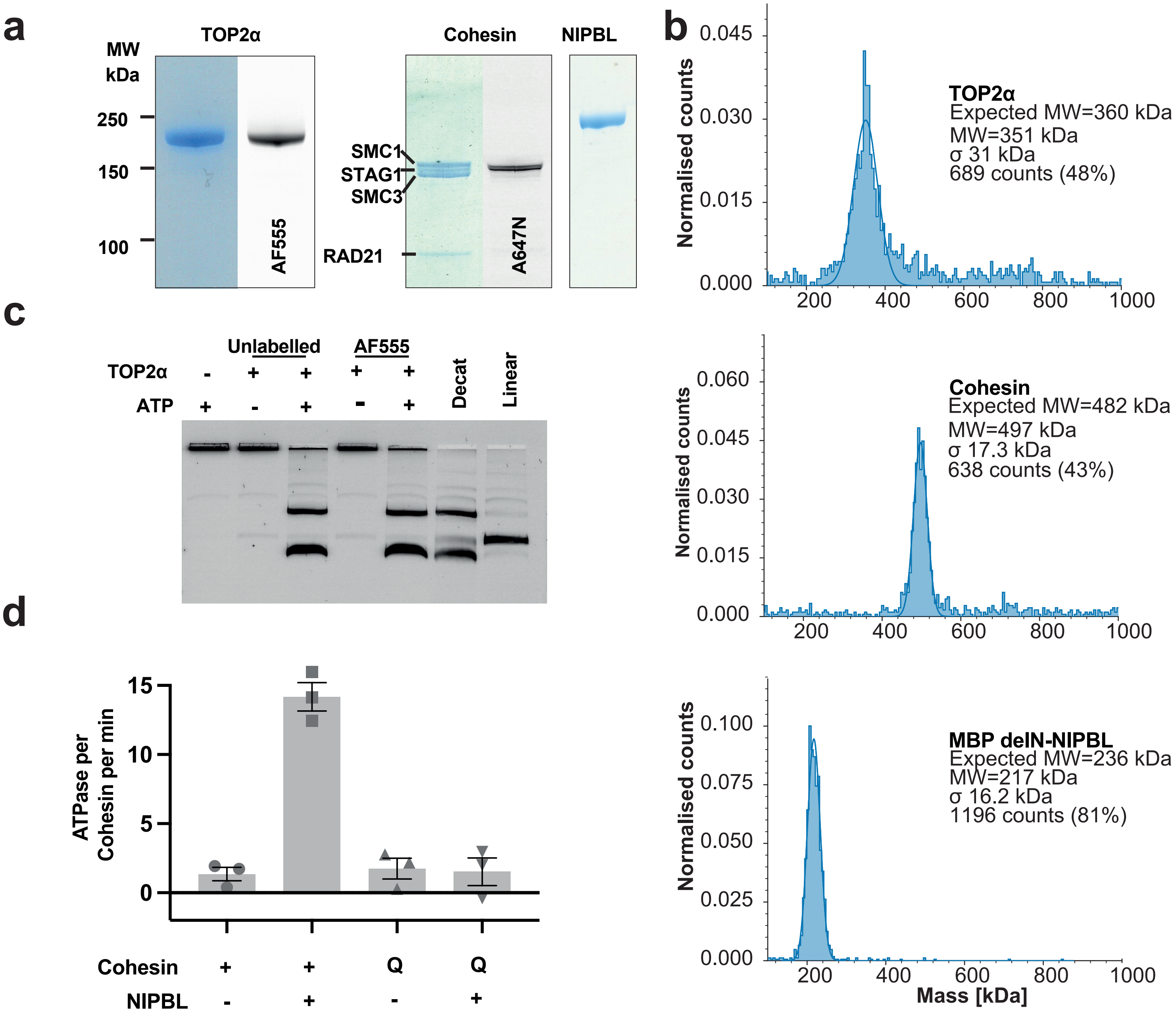


### Supplementary Figure 1

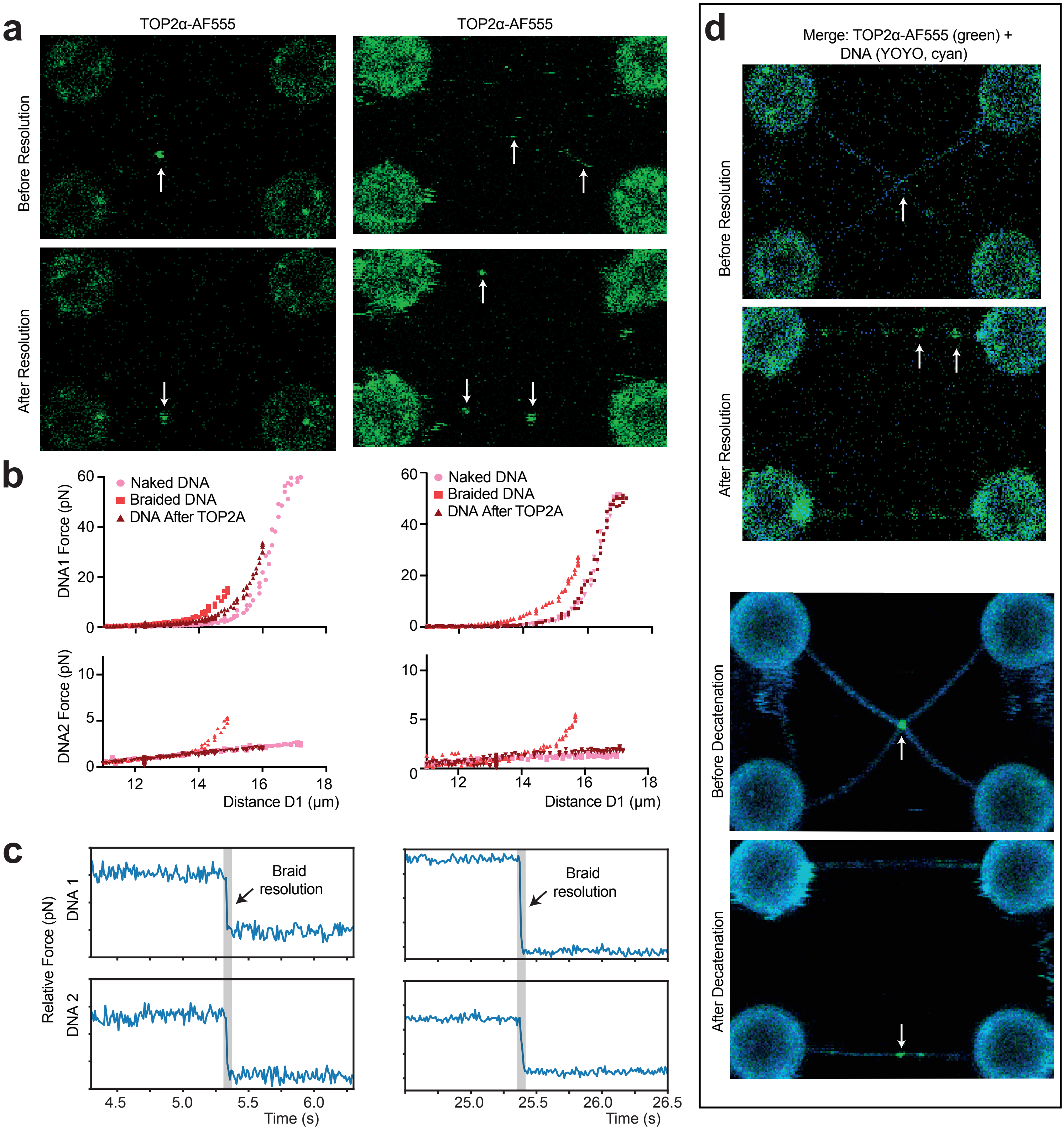


### Supplementary Figure 2

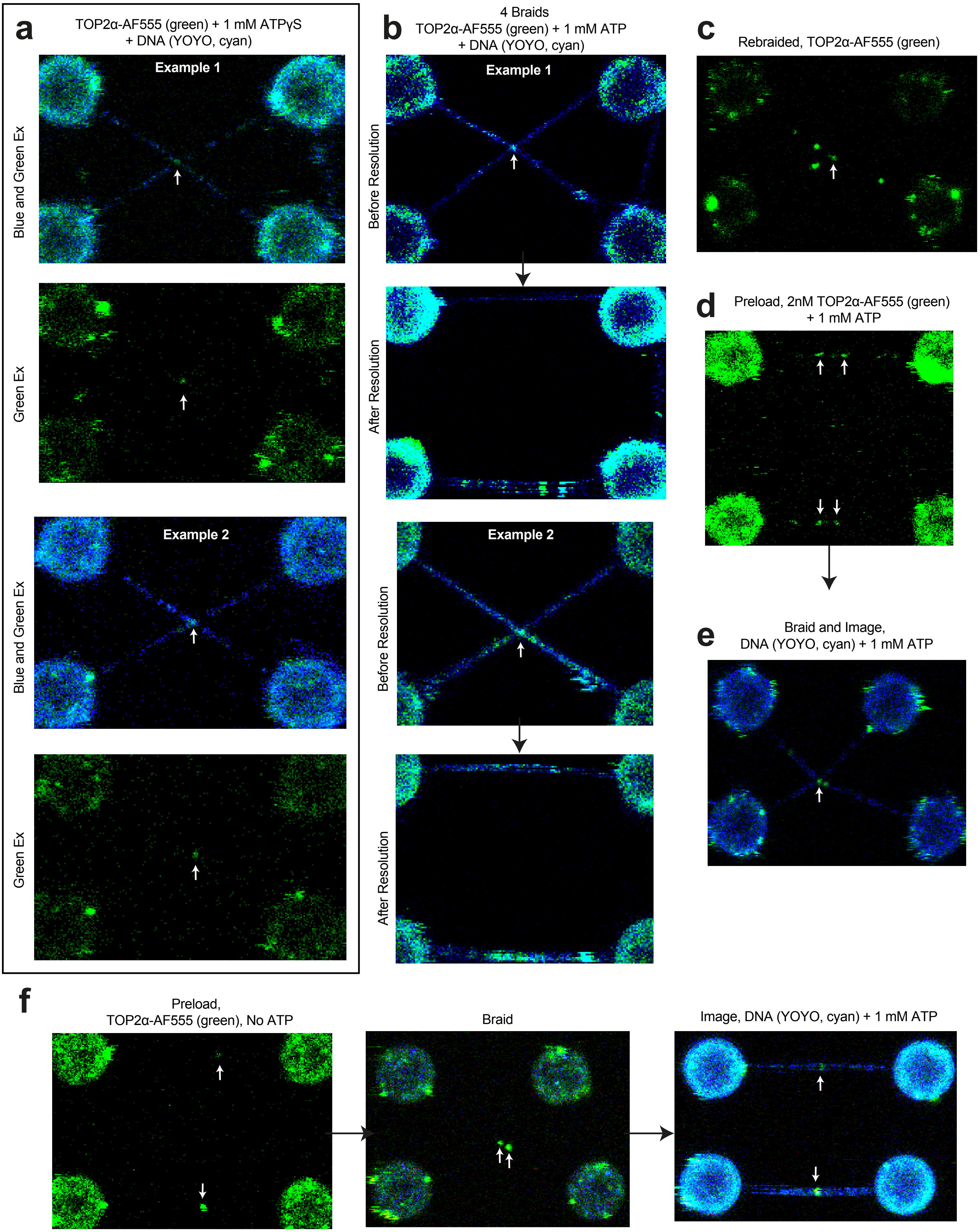


### Supplementary Figure 3
